# Facilitative interaction networks in experimental microbial community dynamics

**DOI:** 10.1101/2023.01.19.524804

**Authors:** Hiroaki Fujita, Masayuki Ushio, Kenta Suzuki, Masato S. Abe, Masato Yamamichi, Yusuke Okazaki, Alberto Canarini, Ibuki Hayashi, Keitaro Fukushima, Shinji Fukuda, E. Toby Kiers, Hirokazu Toju

## Abstract

Understanding potential roles of facilitative interactions between species is one of the major challenges in ecology and microbiology. However, we still have limited knowledge of entangled webs of facilitative interactions in ecosystems. By compiling whole-genome shotgun metagenomic data of an experimental microbial community, we tested the hypothesis that architecture of facilitative interaction networks could change through time. A metabolic modeling approach for estimating dependence between microbial genomes (species) allowed us to infer the network structure of potential facilitative interactions at 13 time points through the 110-day monitoring of experimental microbiomes. We then found that positive feedback loops, which were theoretically predicted to promote cascade breakdown of ecological communities, existed within the inferred networks of metabolic interactions prior to the drastic community-compositional shift observed in the microbiome time-series. We further applied “directed-graph” analyses to pinpoint potential keystone species located at the “upper stream” positions of such feedback loops. These analyses on facilitative interactions will help us understand key mechanisms causing catastrophic shifts in microbial community structure.

## INTRODUCTION

In nature, species form complex webs of interactions, thereby driving various types of community- and ecosystem-level phenomena (May, 1972; Ives and Carpenter, 2007; Allesina and Tang, 2012). Roles of interspecific interactions in sudden shifts of community structure are among the most important targets of ecological research (Scheffer et al., 2001; Scheffer and Carpenter, 2003; Ratzke et al., 2020). Theoretical studies have predicted that architecture (topology) of interaction networks determines consequences of ecological interactions such as species coexistence or community collapse (Thébault and Fontaine, 2010; Rohr et al., 2014; Levine et al., 2017). Although a number of empirical studies on plants and animals have been conducted to test the theories (Olesen et al., 2007; Lever et al., 2014; CaraDonna and Waser, 2020), our knowledge of potential relationship between network architecture and community-level consequences have been limited for microbial ecosystems.

In microbial ecology, estimating architecture of potential interactions itself has been increasingly common (Faust and Raes, 2012; Friedman and Alm, 2012; Berry and Widder, 2014; Kurtz et al., 2015). Amplicon sequencing (DNA metabarcoding) of prokaryote 16S rRNA gene, for example, have been frequently used to infer structure of networks depicting co-occurrence patterns of microbial species (Barberan et al., 2012; Faust et al., 2012; Berry and Widder, 2014). Nonetheless, those networks obtained with co-occurrence pattern analyses include pairs of species that merely share environmental preferences, making it difficult to investigate webs of direct facilitative/competitive interactions between species (Warton et al., 2015; Toju et al., 2017; Kurtz et al., 2019; Blanchet et al., 2020). Moreover, although studies on co-occurrence patterns assume bidirectional associations between species, interspecific interactions in nature are not necessarily bidirectional (Sugihara et al., 2012; Ushio et al., 2018; Delmas et al., 2019). Consequently, reconstructing networks consisting of not only bidirectional but also unidirectional interactions between species (i.e., “directed graphs”) is an essential step for advancing our understanding of microbial community processes.

A promising approach for investigating complex webs of microbial interactions is to estimate flows of metabolites between microbial species based on metagenomic datasets (Stolyar et al., 2007; Klitgord and Segrè, 2010; Zomorrodi and Maranas, 2012; Levy and Borenstein, 2013). Because species’ ability to metabolize given chemical compounds is encoded in their genomes, metabolic modeling has been applied to infer potential metabolic interactions between microbes (Zelezniak et al., 2015; Magnúsdóttir et al., 2017). If genomic information is available for a pair of species, potential dependence of a species on the other species can be evaluated in terms of the list of metabolites presumably emitted by the other species (Zelezniak et al., 2015; Magnúsdóttir et al., 2017). By applying such community-scale metabolic modeling (Frioux et al., 2020), we will be able to gain insights into network architecture of facilitative interactions (Sung et al., 2017; Hassani et al., 2018; Gralka et al., 2020). Analyses on temporal shifts in such metabolic interaction network architecture, in particular, are expected to enhance our understanding of processes or mechanisms causing community collapse (or dysbiosis). Nonetheless, there have been, to our knowledge, no study reporting changes in facilitative interaction network architecture through microbial community dynamics.

In this study, we performed an analysis of metabolic interaction networks using the whole-genome shotgun metagenomic dataset of an experimental bacterial community (Fujita et al., 2022a). Across the 110-day monitoring of a co-culture system of a freshwater bacterial community (Fujita et al., 2022b), the previous study examined temporal shifts in the level of ecological niche overlap between species in order to infer dynamics of competitive interactions (Fujita et al., 2022a). In this study, we reconstructed networks of facilitative interactions based on the metabolic modeling of shotgun metagenomic data at 13 time points across the time-series. We then evaluated changes in the architectural features of the directed graphs through the time-series. Specifically, we tested the hypothesis that positive feedback loops, which have been predicted to destabilize biological communities, existed prior to a sudden community-compositional shift observed in the microbiome experiment. In addition, we examined the presence of microbial species that could be located at the source or sink positions within the directed graphs of metabolic flows. Overall, the preliminary application of community-level metabolic modeling provides a platform for understanding relationship between dynamics of interaction network architecture and ecosystem-level consequences.

## MATERIALS AND METHODS

### Time-series data of the microbial experiment

We used the 110-day time-series dataset of the microbial community experiment described elsewhere (Fujita et al., 2022a, 2022b). In the experiment, the source microbial community was sampled from a pond (“Shoubuike”) near Center for Ecological Research, Kyoto University (34.974 °N, 135.966 °E). The source community was then introduced into the deep-well culture system of an oatmeal broth medium [0.5% (w/v) milled oatmeal (Nisshoku Oats; Nippon Food Manufacturer)] with eight replicates, kept at 23 °C for five days (Fujita et al., 2022b). After the five-day pre-incubation, 200 μL out of 1,000-μL culture medium was sampled from each well of the deep-well plate after mixing (pipetting) every 24 hours for 110 days. In each sampling event, 200 μL of fresh medium was added to each well so that the total culture volume was kept constant. For the samples, amplicon sequencing of 16S rRNA was conducted as reported previously (Fujita et al., 2022b). Based on the amplicon sequencing of the community compositional dynamics, we selected the replicate community that showed the largest community compositional changes within the time-series (Fujita et al., 2022a): a rapid and substantial community compositional change occurred around Day 18 in the replicate community (Fig. 1). The extracted DNA samples of the replicate community was subjected to a whole-genome shotgun sequencing analysis, which targeted 13 time points across the time-series (Day 1, 10, 20, 24, 30, 40, 50, 60, 70, 80, 90, 100, 110; ca. 10 Gb per sample) (Fujita et al., 2022a). The analysis described below was performed by compiling the whole-genome shotgun sequencing data (Fujita et al., 2022a).

**FIGURE 1.**
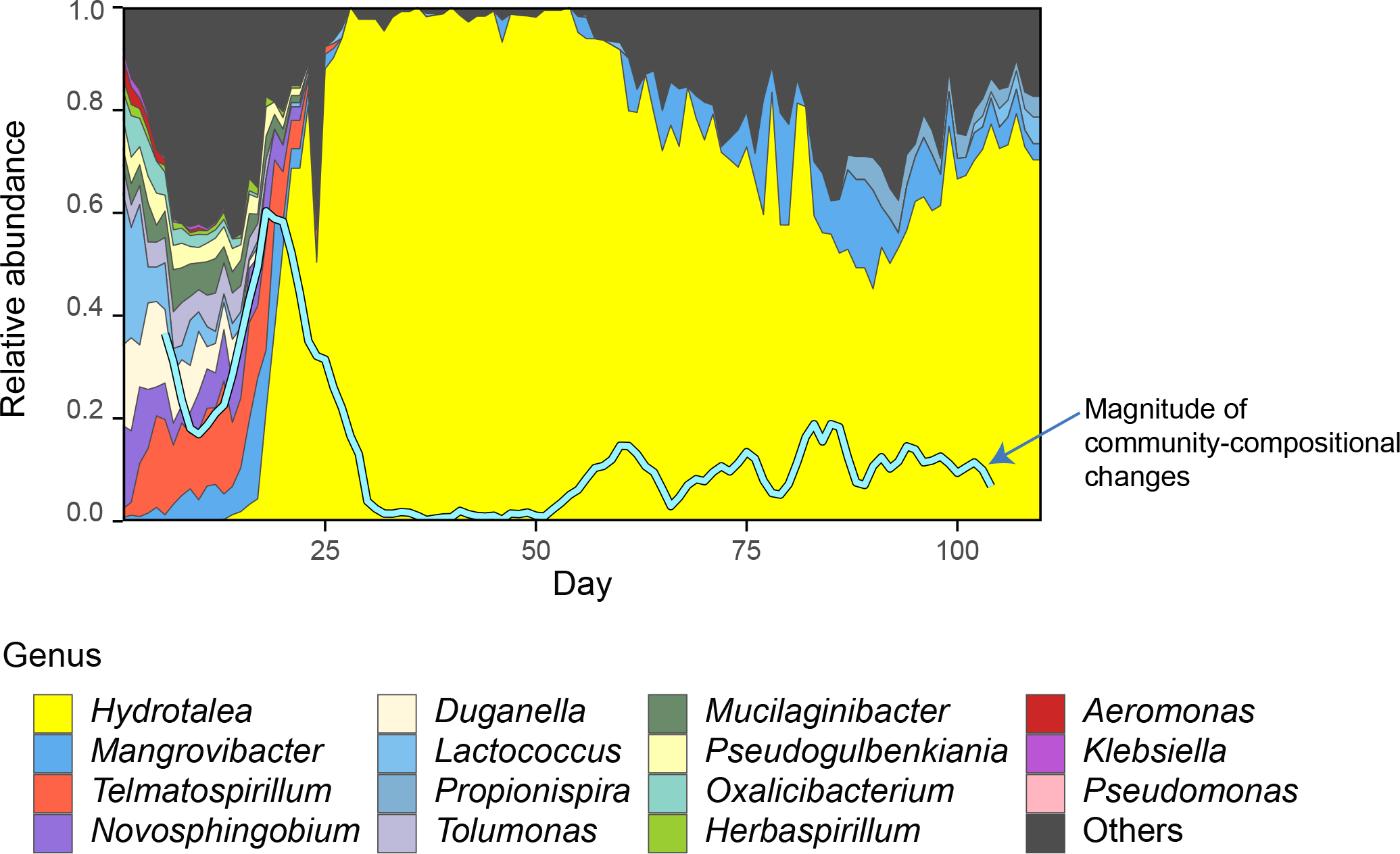
Time-series data of the community structure. Through the 110-day experiment, community compositions were monitored based on 16S rRNA sequencing. To quantify the speed and magnitude of community shifts through time, the “abruptness” index was calculated through the time-series (blue line). Specifically, an estimate of the abruptness index for time point *t* was obtained as the Bray-Curtis *β*-diversity between average community compositions from time points *t* – 4 to *t* and those from *t* + 1 to *t* + 5 (i.e., dissimilarity between 5-day time-windows). An abruptness score larger than 0.5 indicates that turnover of more than 50 % of community compositions occurred between the time-windows. Reproduced from the amplicon sequencing data of a previous study on the microbiome system (Fujita *et al.*, 2022b).

### Whole-genome shotgun metagenomics

The whole-genome shotgun sequencing data were processed as detailed previously (Fujita et al., 2022a). Briefly, after adaptor trimming with Cutadapt (Martin, 2011) and quality filtering with Fastp0.21.0 (Chen et al., 2018) [in total, the number of output sequencing reads was 1002.49 M (160.08 Gb)], the sequences of each time-point sample were assembled using metaSPAdes 3.15.2 (Bankevich et al., 2012). Binning and quality assessing, were then performed with MetaWRAP 1.3.2 (Uritskiy et al., 2018) and CheckM 1.1.3 (Parks et al., 2015), respectively. The identity between MAGs were calculated with FastANI 1.33 (Jain et al., 2018) and MAGs with > 99 % identity were grouped through the time-series. The read-coverage calculation was then conducted using CoverM 0.6.0 (Woodcroft B, 2021).

Taxonomic annotation and genome annotation were conducted, respectively, with GTDB-Tk 1.6 (Chaumeil et al., 2020; Parks et al., 2022) and Prokka 1.14.6 (Seemann, 2014) 1.14.6. The metagenome-assembled genomes (MAGs) with > 80 % completeness and < 5 % contamination were used in the analyses below. The orthology numbers of Kyoto Encyclopedia of Genomes (KEGG) were retrieved for respective genes using GhostKOALA 2.2 (Kanehisa et al., 2016) and the completeness of metabolic pathways was estimated for each MAG using KEGG decoder 1.3 (Graham et al., 2018). In total, 32 MAGs belonging to 20 genera (16 families; 12 orders) were detected across the time-series (Supplementary Data 1) (Fujita et al., 2022a).

### Metabolic modeling

To explore potential effects of facilitative interactions between microbes within the microbiome, we performed an analyses of metabolite-exchange interaction networks based on the MAGs detailed above. For each MAG, we reconstructed a metabolic model based on the top-down carving approach of curated “universal models” (i.e., manually curated and simulation-ready metabolic models) (Machado et al., 2018) using CarveMe 1.5.0 (Machado et al., 2018). Potential metabolic interactions between microbial MAGs were then evaluated based on species coupling scores indicating dependency of target species in the presence of others as implemented in SMETANA 1.0.0 (Zelezniak et al., 2015). In this approach, all potential exchanges of metabolites between species were mapped with the default parameters as implemented in SMETANA.

### Network analysis

The inferred metabolic interaction network of each time point was then analyzed based on the treeness, feedforwardness, and orderability (Corominas-Murtra et al., 2013) Treeness is a measure of pyramidal (top-down) network structure, in which small numbers of nodes at upper layers have outward links to many other nodes at lower layers. Feedforwardness is a measure of network-scale bias in the direction of links: a high feedforwardness value represents strong upstream-downstream structure within a network. Meanwhile, orderability represents the degree of the lack of feedback loops within directed graphs (networks). As the orderability index is defined as the proportion of nodes outside feedback network loops, it ranges from 0 (loop structure involving all nodes) to 1 (absence of loops).

To evaluate topological positions of respective microbial MAGs within the networks, influence (Masuda et al., 2009) (a measure of the degree to which a focal node has influence on the others within a directed graph) and PageRank centrality (Page and Brin, 1998) (a measure of the degree to which a focal node has links from other nodes with many inward links) were calculated.

## RESULTS

### Ecosystem-level profiles

After the drastic change in community structure around Day 18 (Fig. 1), the community-level compositions of metabolic pathways/processes greatly changed (Fig. 2). For example, the function of sulfite dioxygenase and that of NO_2_^−^/NO/N_2_O reduction pathways seemed to decline by Day 30 and 40, respectively (Fig. 2). Although microbes encoding these functions in their genomes might still exist at a small proportion (under detection limit of our sequencing analysis), rapid alternations of major functional profiles presumably occurred in the microbial ecosystem through the time-series.

**FIGURE 2.**
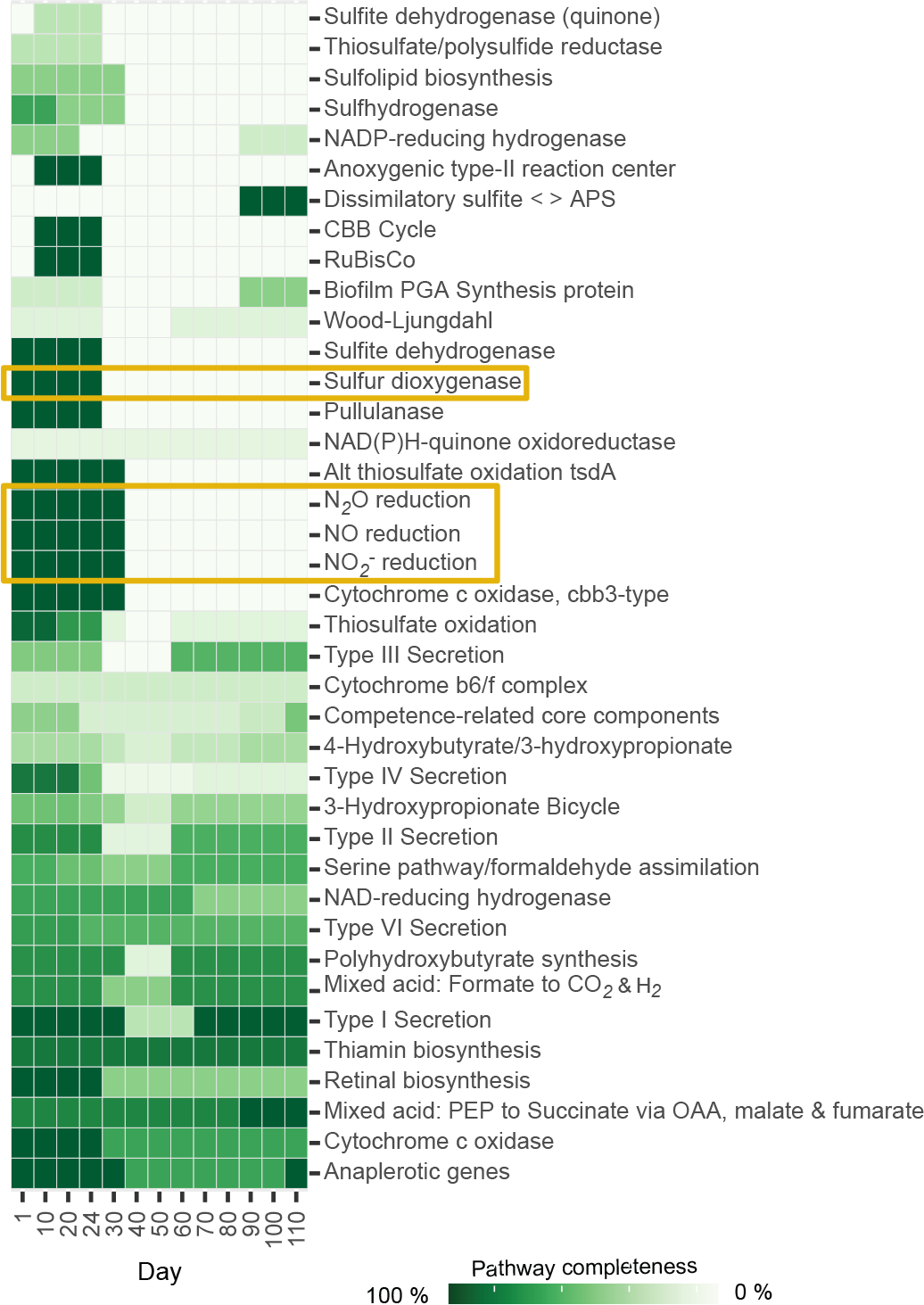
Highlights of changes in community-level profiles of metabolic pathways/processes through the time-series. After assembling the data of all the MAGs detected on each day, community-level pathway completeness is shown for the pathways/processes that exhibited temporal changes in pathway completeness. To focus on the metabolic pathways/processes that varied greatly through time, the pathways/processes whose metagenome-level completeness exceeded 0.9 at 12 or more time points are not shown. Metabolic pathway/process profiles mentioned in the main text are highlighted.

### Dynamics of metabolic interaction networks

Microbial MAGs belonging to different taxa were linked with each other within the network of potential facilitative interactions (Fig. 3; Supplementary Data 2). In particular, microbes in the class Gammaproteobacteria were inferred to provide metabolites to microbes in other taxonomic groups. Likewise, *Terracidiphilus* bacteria (Acidobacteriae) had links of potential metabolite supply towards some bacteria belonging to Gammaproteobacteria and Alphaproteobacteria at some time points (Fig. 3). The number of detectable nodes suddenly decreased between Days 20 and 30, entailing rapid decline of the inferred metabolic interaction networks (Fig. 3). The microbial community then reached a quasi-stable state characterized by several bacteria in the genera *Hydrotalea*, *Terracidiphilus*, *Mangrovibacter*, and *Rhizomicrobium* (from Day 40 to Day 50; Fig. 3). Among them, unidirectional facilitative effects from *Mangrovibacter* to other bacteria were inferred based on the metabolic modeling analysis (Fig. 3). The number of detectable MAGs gradually increased from Day 60, resulting in the restoration of an entangled web of potential metabolic interactions on Day 110 (Fig. 3).

**FIGURE 3.**
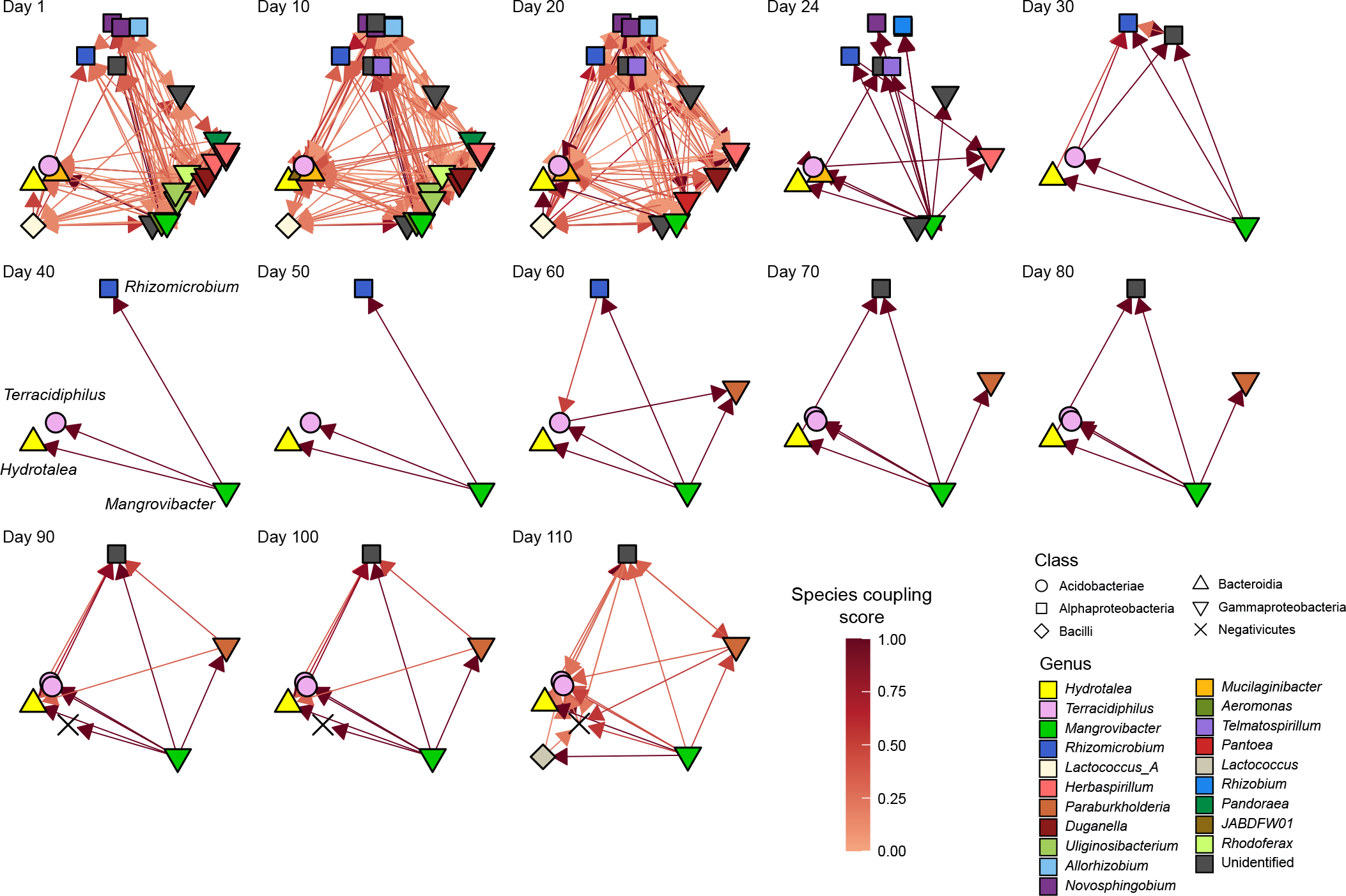
Inferred network of metabolic interactions between microbes. Based on the whole-genome shotgun metagenomic data, genome-scale metabolic modeling was conducted at each of the target time point. The results were used to infer potential flows metabolites between microbial MAGs. Positive effects inferred by metabolic modeling are shown with arrows connecting donor and recipient microbial MAGs. Darker colors of arrows indicate higher species coupling scores inferred in the metabolic modeling analysis.

The treeness, feedforwardness, and orderability of the network of the potential metabolic interactions varied considerably across the time-series (Fig. 4). Until Day 20, the network structure was characterized by low treeness, low feedforwardness, and low to moderate orderability (Fig. 4b). The facilitative interaction network then showed drastic architectural shift until Day 40 as characterized by the rapid increase of orderability (Fig. 4b). This result indicates that the dynamics of the network architecture are characterized by the presence of positive feedback loops (as represented by low orderability) early in the time-series and that such feedback loops disappeared from the microbial community by Day 40 (Fig. 3). Through the gradual restoration of network complexity after Day 60, the presence of feedback loops was inferred again on Day 110 (Fig. 3) as indicated by lowered network orderability estimate on the day (Fig. 4).

**FIGURE 4.**
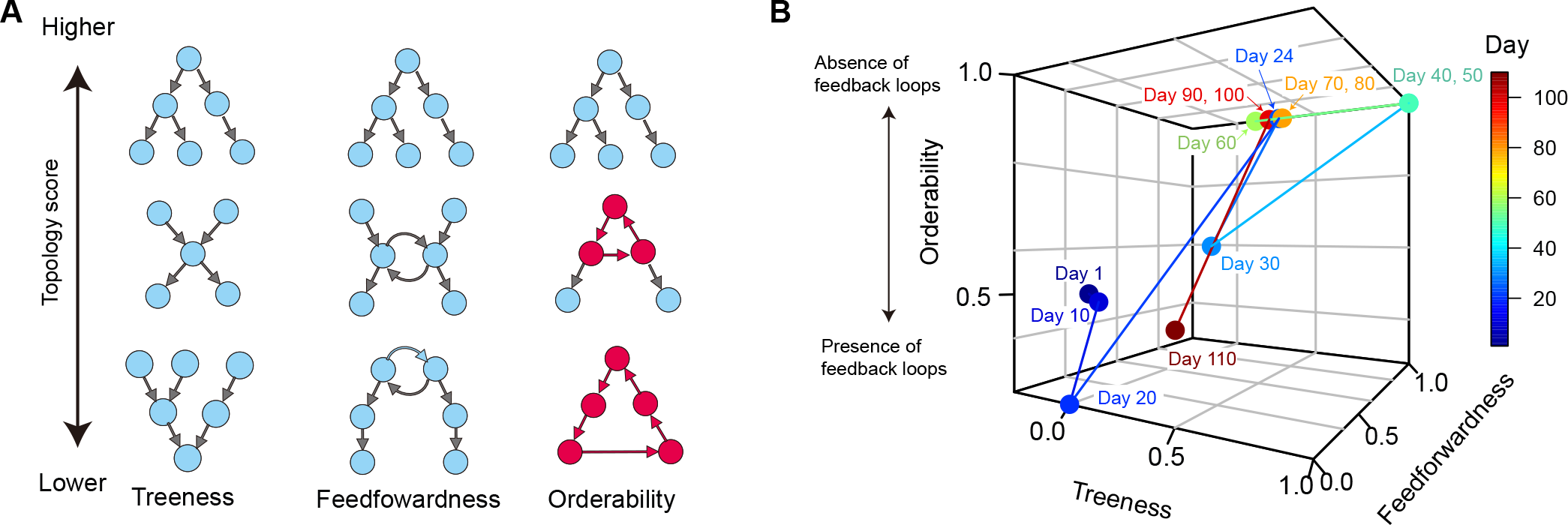
Network topology analysis. **(A)** Schema of network architectural properties. Treeness and feedforwardness represent pyramidal and upstream-downstream structures of directed graphs, respectively. Orderability represents lack of feedback loops within a network. Along the axis of orderability, the nodes and links included in feedback loops are highlighted in red. **(B)** Dynamics of network characteristics. Changes in network architectural properties are shown in terms of treeness, feedforwardness, and orderability. Networks with low “orderability”, by definition, contain loops of flow of metabolites, while those with maximum orderability (= 1) lack feedback loops.

### Potential keystone species

Within the metabolic interaction networks (Fig. 3), some microbial MAGs belonging to the class Gammaproteobacteria were located at the “upper stream” of the network, showing high influence scores (Fig. 5; Supplementary Data 1). In particular, a gammaproteobacterial MAG in the genus *Mangrovibacter* consistently showed the highest influence among the microbes detected at most time points (Fig. 5). Meanwhile, microbes located at the sink positions within the inferred metabolic interaction networks (i.e., MAGs with high PageRank scores) represented diverse taxonomic groups (Fig. 5). From Day 40 to 50, through which a small number of bacterial taxa represented the microbiome structure, simple source–sink relationship of potential metabolite flow was observed between *Mangrovibacter* and others (i.e., *Hydrotalea*, *Terracidiphilus*, and *Rhizomicrobium*; Figs. 4 and 5).

**FIGURE 5.**
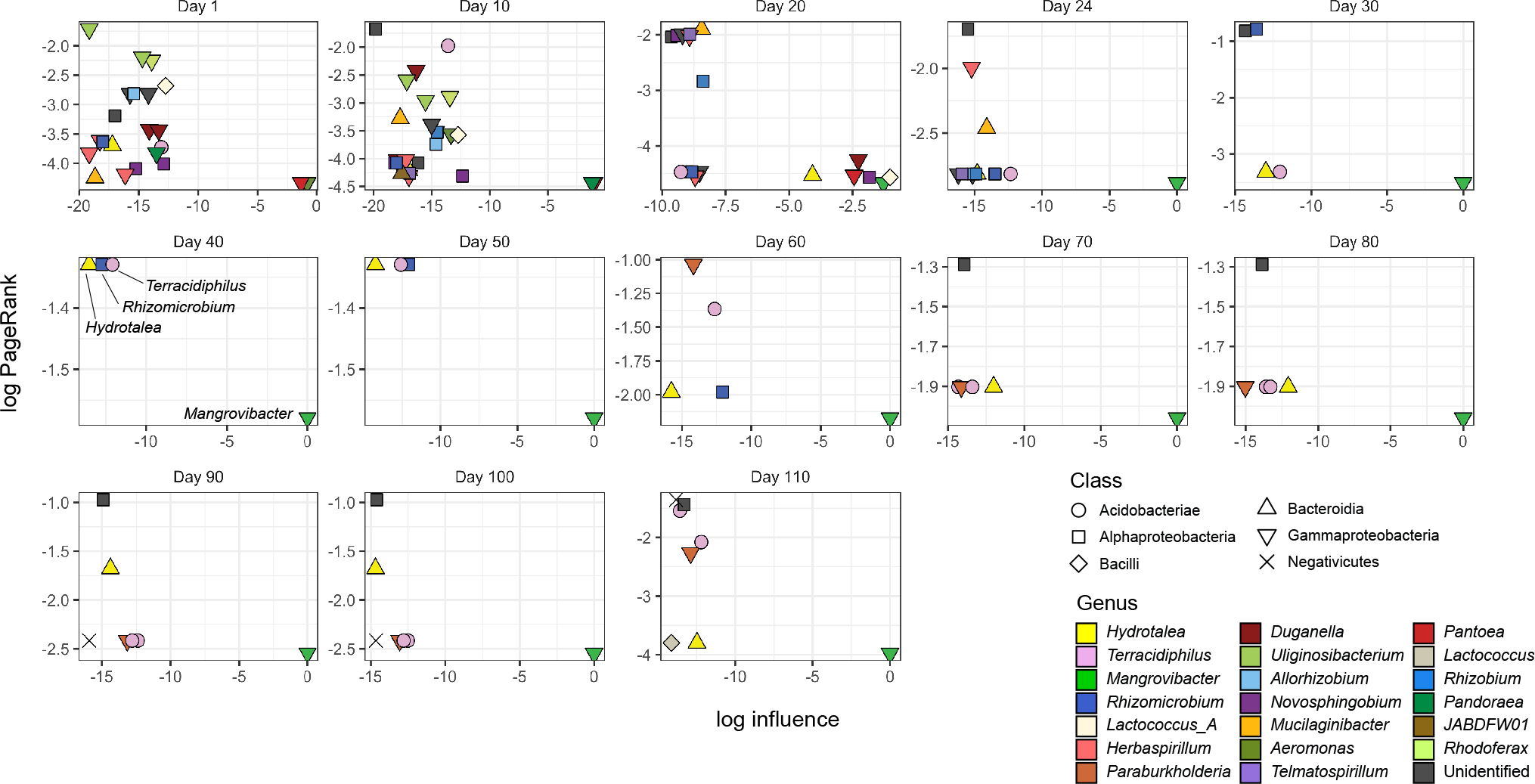
Potential keystone species/taxa within metabolic interaction networks. Within each directed graph of metabolic dependence network (Fig. 4), influence (a measure of the degree to which a focal node has influence on the others within a directed graph) and PageRank (a measure of the degree to which a focal node has links from other nodes with many inward links) measures of network centrality was calculated for each microbe. *Mangrovibacter* tended to show high impacts (influence) on other bacteria within the metabolic interaction networks throughout the time-series.

## DISCUSSION

We showed preliminary results on temporal shifts in the network architecture of facilitative interactions by compiling a whole-genome shotgun metagenomic dataset of experimental microbiome dynamics. While ecosystem-level profiles of metabolic functions (Fig. 2) have been intensively investigated (Raes and Bork, 2008), shotgun metagenomic data also allow us to infer ecological processes of species interactions (Figs. 3–5). Classic theory predicts that facilitative interactions basically destabilize biological communities (Allesina and Tang, 2012). However, recent theoretical studies suggest that such effects of facilitative interactions depend on network architecture of interactions (Bastolla et al., 2009; Thébault and Fontaine, 2010; Fontaine et al., 2011; Morton et al., 2022). Nestedness, for example, have been intensively investigated as a potential key property of facilitative interaction networks in terms of species coexistence (Bascompte et al., 2003; Thébault and Fontaine, 2010; Rohr et al., 2014). Meanwhile, most studies on facilitative ecological interactions have relied on the assumption that all links within a network are bidirectional (i.e., mutualistic). In this study, we explored ways for uncovering the structure of directed graphs of species interactions (Sugihara et al., 2012; Ushio et al., 2018; Delmas et al., 2019) based on a metagenomic analysis of potential metabolic interactions (Zelezniak et al., 2015). Our finding that architecture of directed interaction networks could drastically change through time will fuel discussion on potential roles of interaction network structure on biological community dynamics and stability.

Among the directed-graph indices examined in this study, orderability was of particular interest (Fig. 4). It has been theoretically predicted that presence of positive feedback loops in facilitative interaction networks can destabilize ecological communities (Coyte et al., 2015; Levine et al., 2017). Specifically, such feedback structure of dependence may magnify cascades of population collapse once balance of population size among constituent species fluctuates within the feedback loops. In a previous study on the examined experimental microbiome, a high level of niche overlap among bacterial species was inferred to have promoted community compositional shifts (Fujita et al., 2022a). In particular, niche overlap within the gammaproteobacterial or alphaproteobacterial sub-community (guild) presumably resulted in competitive exclusion of constituent microbial populations (Fujita et al., 2022a). Such competition-driven decline of some gammaproteobacterial or alphaproteobacterial species may have triggered a cascade breakdown of species (Rezende et al., 2007) through the positive feedback loop observed in this study (Fig. 3). In other words, once competitive exclusion occurs within an ecological guild, species depending on the metabolites of the declining guilds are expected to be negatively influenced by the reduced flow of metabolites through the facilitative interaction network.

Treeness and feedforwardness of network architecture give additional important information about propagation of negative effects within networks. If a facilitative interaction network has hierarchical structure represented by high treeness and feedforwardness (Corominas-Murtra et al., 2013), placement of the ecological guilds from which fluctuations are initiated would influence subsequent ecological processes through the network. Specifically, fluctuations occurred in upstream positions may be propagated more rapidly throughout the network, while those derived from downstream positions would entail minimal impacts on the entire community. Albeit the potential roles of such hierarchical structure, the network architecture observed early in the experimental community (until Day 20) was represented by low treeness and feedforwardness (Fig. 4b). Thus, influence of hierarchical network structure on community collapse remains to be examined in future studies on networks with high treeness and feedforwardness.

In parallel with investigations on the entire network structure, directed-graphs reconstructed with metagenomic data provide us with insights into species occupying upstream/downstream positions within networks. Species located at upstream positions within a “supply chain” of metabolites may impose greater impacts on population dynamics of other species within the network than species at downstream positions. In our data, a bacterium in the genus *Mangrovibacter* continued to occupy upstream positions throughout the community dynamics as indicated by the analysis of network influence scores (Fig. 5). Thus, although the *Mangrovibacter* bacterium was a minor component of the community (Fig. 1), it might have disproportionately large impacts on the dynamics of the entire microbiome. The working hypothesis can be tested by removing the *Mangrovibacter* bacterium from the experimental system. Nonetheless, such selective removal of specific bacterial species from microbiomes remains a challenge because the use of antibiotics often causes unexpected side-effects on non-target species (Cho et al., 2012; Francino, 2016; Langdon et al., 2016). Technical advances that allow selective removal of potential “keystone species” (Paine, 1966; Power et al., 1996) within microbiomes are awaited.

Beyond the preliminary results obtained in this study, further studies based on metabolic modeling approaches are required to understand dynamics and consequences of facilitative interactions in ecological communities. Context-dependency of network architecture, for example, needs to be examined by comparing network dynamics among different experimental settings (e.g., different culture media or different temperature conditions) (Zelezniak et al., 2015; Magnúsdóttir et al., 2017). It is also important to evaluate to what extent network architectural properties inferred with the metabolic modeling approaches are consistent with those estimated with other informatics approaches. In this respect, comparison with recently developed methods for reconstructing species interactions based on time-series data is of particular interest (Deyle et al., 2016; Ushio et al., 2018; Suzuki et al., 2022). Furthermore, integrating information of facilitative interactions with that of competitive interactions is an essential step for examining how relative balance of multiple interaction types affect community stability (Bastolla et al., 2009; Fontaine et al., 2011; Mougi and Kondoh, 2012; Goldford et al., 2018). Interdisciplinary studies combining genomics and ecological theory will broaden our views on fundamental mechanism driving microbial community dynamics.

## Data availability statement

The datasets and codes used in this study can be found in online repositories. The names of the repository/repositories and accession number(s) can be found below:

Github, https://github.com/hiroakif93/Facilitative-interaction-networks-in-experimental-microbial-community-dynamics-DDBJDRA, https://www.ddbj.nig.ac.jp/, DRA013382.

## Author contributions

HT designed the work with HF. HF performed experiments. HF analyzed the data with HT. HF and HT wrote the paper with all the authors.

## Acknowledgements

We thank Sayaka S. Suzuki and Keisuke Koba for support in the experiment.

